# Chemical inhibition of RPA gap protection sensitizes BRCA1-deficient cancers to PARP inhibition

**DOI:** 10.1101/2025.04.09.647957

**Authors:** Pamela S. VanderVere-Carozza, Matthew R Jordan, Joy E. Garrett, Karen E. Pollok, Hilary D. Hinshaw, Xierzhatijiang Sulaiman, Sheng Liu, Jun Wan, Katherine S. Pawelczak, John J. Turchi

**Affiliations:** Department of Medicine, Indiana University School of Medicine; Indianapolis, IN, United States; Department of Radiation Oncology, Indiana University School of Medicine; Indianapolis, IN, United States; Herman B. Wells Center for Pediatric Research, Departments of Pediatrics, Pharmacology and Toxicology, Medical and Molecular Genetics Indiana University; Indianapolis, IN, United States; Indiana University Cancer Center, Indiana University School of Medicine; Indianapolis, IN, United States; Department of Obstetrics and Gynecology, Division of Gynecologic Oncology, Indiana University School of Medicine; Indianapolis, IN, United States; Department of Bioinformatics, Indiana University School of Medicine; Indianapolis, IN, United States; NERx BioSciences; Indianapolis, IN, United States

**Author notes:** These authors contributed equally (Pamela S. VanderVere-Carozza, Matthew R Jordan).

## Abstract

Poly (ADP-ribose) polymerase inhibitors (PARPi) are standard of care for many BRCA1 deficient cancers, though few cures are achieved. We sought to determine if targeting the protection of the single-strand DNA gaps induced by PARPi in BRCA-deficient cancers could increase efficacy. Replication protein A (RPA) participates in critical protein-protein and protein-DNA interactions to protect single-stranded DNA (ssDNA) and support DNA metabolism. We have reported the optimization of small molecule RPA inhibitors (RPAi) that target protein-ssDNA interactions to chemically exhaust RPA and elicit single-agent anticancer activity. RPAi sensitizes cells to PARPi in BRCA1-deficient non-cancerous cells, where ssDNA gap formation drives therapeutic efficacy. We show that RPAi treatment abolishes PARPi-induced replication gap protection, resulting in genomic instability via replication fork degradation and chromosomal integrity, and that, *in vivo*, the RPA and PARP targeted combination abrogates cancer growth in a BRCA1-mutant breast cancer model. We find that genetic predispositions to ssDNA gap accumulation correlate with RPAi sensitivity, and BRCA1-proficient cells remain sensitive to combination treatment but require more PARP inhibition to increase ssDNA gaps. RPAi-PARPi combination activity in patient-derived ovarian cancer models demonstrates the utility of targeting gap protection to increase PARPi sensitivity and circumvent resistance. Collectively, this work provides a unifying mechanism of chemical RPA exhaustion as a cancer therapeutic strategy.

**One Sentence Summary:** Inhibition of single-strand DNA gap protection potentiates PARP targeted treatment of BRCA1 deficient cancers.

## INTRODUCTION

Recent advances in our understanding of DNA damage response (DDR) signaling have opened up an array of opportunities to treat human cancer [1]. The DDR is initiated by the phosphatidylinositol 3-kinase-related kinases (PIKKs) ATM, ATR, and DNA-PK, each of which responds to different DNA structures [2]. The PIKKs each require a sensor protein to bind to DNA: Ku binds to double-stranded DNA (dsDNA) ends to activate DNA-PK, MRN also binds to double-strand breaks to activate ATM, and replication protein A (RPA) binds to single-stranded DNA (ssDNA) to activate the ATR kinase. Once activated, these kinases signal to downstream kinases and other effector proteins to regulate the cell cycle and DNA replication and promote DNA repair, and if the damage is not repaired, cell death via apoptosis or mitotic catastrophe ensues. RPA, a human ssDNA-binding protein, is a crucial sensor of ssDNA resulting from replication stress (RS)[3]. The binding of the ATR-interacting protein (ATRIP) to the RPA-ssDNA complex results in the activation of ATR and initiation of the DDR [4, 5], limiting replication origin firing and inducing cell cycle arrest to maintain genome stability. While the majority of DDR-targeted anticancer therapeutics are directed toward transducer PIKKs and downstream effector kinases, we have developed a small-molecule RPA inhibitor (RPAi), NERx-329, that blocks the interaction between the RPA sensor ssDNA [6]. RPAi treatment has broad single-agent anticancer activity and synergizes with DNA-damaging agents and other DDR inhibitors [7], including poly (ADP-ribose) polymerase inhibitors (PARPi).

PARPi therapy is approved for treating a variety of cancers that harbor genetic defects in homologous recombination (HR) via a synthetic lethal interaction[8]. Recent data showing that BRCA1 deficiency results in the formation of lagging strand gaps that can be exacerbated by the inhibition of PARP indicate that lagging strand ssDNA gaps are an important determinant of PARPi sensitivity [9]. Within this context, the ssDNA gaps generated in the initial S phase persist through the cell cycle and become DSBs during replication in the next S phase, thereby resulting in replication fork collapse [10, 11]. Moreover, in BRCA1-deficient cells with acquired PARPi resistance, excessive ssDNA gaps are still generated through nucleolytic nick/SSB expansion independent of PARPi treatment [12]. As both PARP proteins and BRCA1 participate in gap suppression, the sensitivity of BRCA1-deficient cells to PARPi is, in part, a function of dysregulated lagging strand replication[9, 13]. We have demonstrated that RPA function is critical to the protection of these gaps and that chemical inhibition of RPA increases the sensitivity of BRCA1-deficient RPE cells to PARP inhibition [9]. Moreover, the proposal that ssDNA gaps are the DNA lesions that drive the therapeutic response [14] has stimulated recent investigations into many other processes that increase ssDNA gap formation [15-18], and RPA inhibition is also predicted to trigger sensitivity.

Consistent with the importance of RPA in lagging strand gap suppression, the RPA protection threshold is established by the expression and activity of RPA to bind and protect ssDNA[3, 19]. Under normal physiological conditions, the amount of ssDNA generated remains below the RPA protection capacity. RS induced by oncogene activation results in MCM loading defects, premature origin complex assembly, and RPA-dependent origin firing [20], in which BRCA1 deficiency generates lagging strand gaps. In both cases, the amount of single-stranded DNA remaining below the RPA protection threshold; hence, the cells can continue growing and dividing. Replication stress induced by DNA damage, replication fork stalling, or chemical inhibition of the DDR can elevate the ssDNA level above the RPA protection threshold, resulting in replication catastrophe, which accounts for the cell death observed following these events [19, 20]. This analysis is significant in that it explains the activity of common chemotherapeutics and their therapeutic window in treating cancers but also explains the activity of DDR-targeted therapies [3]. Therefore, the therapeutic window of RPA inhibition requires sufficient RPA to protect ssDNA under normal physiological conditions but insufficient RPA in cells with oncogenic RS to achieve single-agent anticancer activity. Additionally, this should result in even greater efficacy when combined with DNA-damaging or RS-inducing treatments. The ability to increase the effectiveness of DNA-damaging therapeutics or targeted DDR inhibitors can thus be achieved by either generating more ssDNA to allow the state of RPA exhaustion to be achieved or by reducing the level of active RPA (chemical RPA exhaustion) such that less single-stranded DNA is needed to achieve RPA exhaustion and replication catastrophe [21-23]. In this work, we elucidate the mechanistic impact of RPA exhaustion in the context of ssDNA gaps. Chemical RPA inhibition results in the degradation of lagging strand gaps in BRCA1-deficient cells, which is exacerbated by PARP inhibition, leading to chromosome pulverization and cell death. Moreover, this translates to the *in vivo* efficacy of the RPAi-PARPi combination treatment. RPA exhaustion therefore has the potential to provide a unifying mechanism related to the efficacy of not only DNA-damaging therapeutics but also PARP inhibition sensitivity and resistance, suggesting that an RPA inhibitor will provide effective therapeutic options in a variety of different tumor types.

## RESULTS

### The RPAi NERx-329 blocks protection of PARPi-induced gaps

We previously demonstrated in a noncancer retinal pigment epithelial (RPE) BRCA1 knockout cell culture model that chemical RPA inhibition enhanced PARPi (olaparib)-induced cell death [9]. To assess the underlying mechanism of this potential RPAi-PARPi therapeutic combination, we employed a BRCA1-deficient MDA-MB-436 triple-negative breast cancer (TNBC) model and DNA fiber combing with S1 nuclease to determine the impact on replication fork dynamics and ssDNA gap formation[27, 28]. After 2 hours of individual or concurrent combination treatment, replication fork progression was monitored by the incorporation of the thymidine analog 5-chloro-2’-deoxyuridine (CldU) into DNA, which was then detected by fluorescently labeled antibodies (Figure 1A). Within each replicate, the collected cells were split into two fractions and processed identically to treat the released DNA with and without S1 nuclease immediately before being combed onto coverslips. S1 nuclease nicks ssDNA; thus, fiber length shortening after S1 treatment is indicative of the presence of ssDNA gaps [29]. Consistent with BRCA1-deficient gap formation [30], control cultures treated with vehicle revealed the presence of ssDNA gaps, as evidenced by the decrease in fiber length upon fiber processing with S1 nuclease (Figure 1A). Treatment with NERx-329 did not significantly affect fiber length, and ssDNA gaps were still observed, suggesting that NERx-329 treatment does not completely exhaust RPA under these experimental conditions (Figure 1A). Cells treated with olaparib exhibited accelerated replication and significantly increased fiber length derived from extensive ssDNA gap formation [9, 31], as expected (Figure 1A). Strikingly, the combination of NERx-329 with olaparib reversed the apparent acceleration of replication and resulted in significantly shortened fibers devoid of ssDNA gaps (Figure 1A). These data could be a result of DNA defects in replication, as we observed in BRCA wild-type NSCLC A549 cells, which exhibited slower replication after pretreatment with NERx-329 (Figure S1A) or degradation of ssDNA gaps from the loss of protection due to chemical RPA exhaustion. To delineate these two possibilities, we included the MRE11 inhibitor mirin in the treatments with olaparib alone and in combination with NERx-329. Importantly, MRE11 is a critical nuclease responsible for degrading replication fork-dependent ssDNA gaps [32]. Similar to treatment with olaparib alone, treatment with olaparib or mirin resulted in the formation of longer fibers still containing ssDNA gaps (Figure 1A). Importantly, the addition of mirin to MDA-MB-436 cells treated with olaparib and NERx-329 resulted in longer DNA fibers than the combination without mirin did and still resulted in ssDNA gaps, although the fiber lengths were not completely rescued to the length of the olaparib and mirin treatment conditions (Figure 1A).

**Fig. 1.**
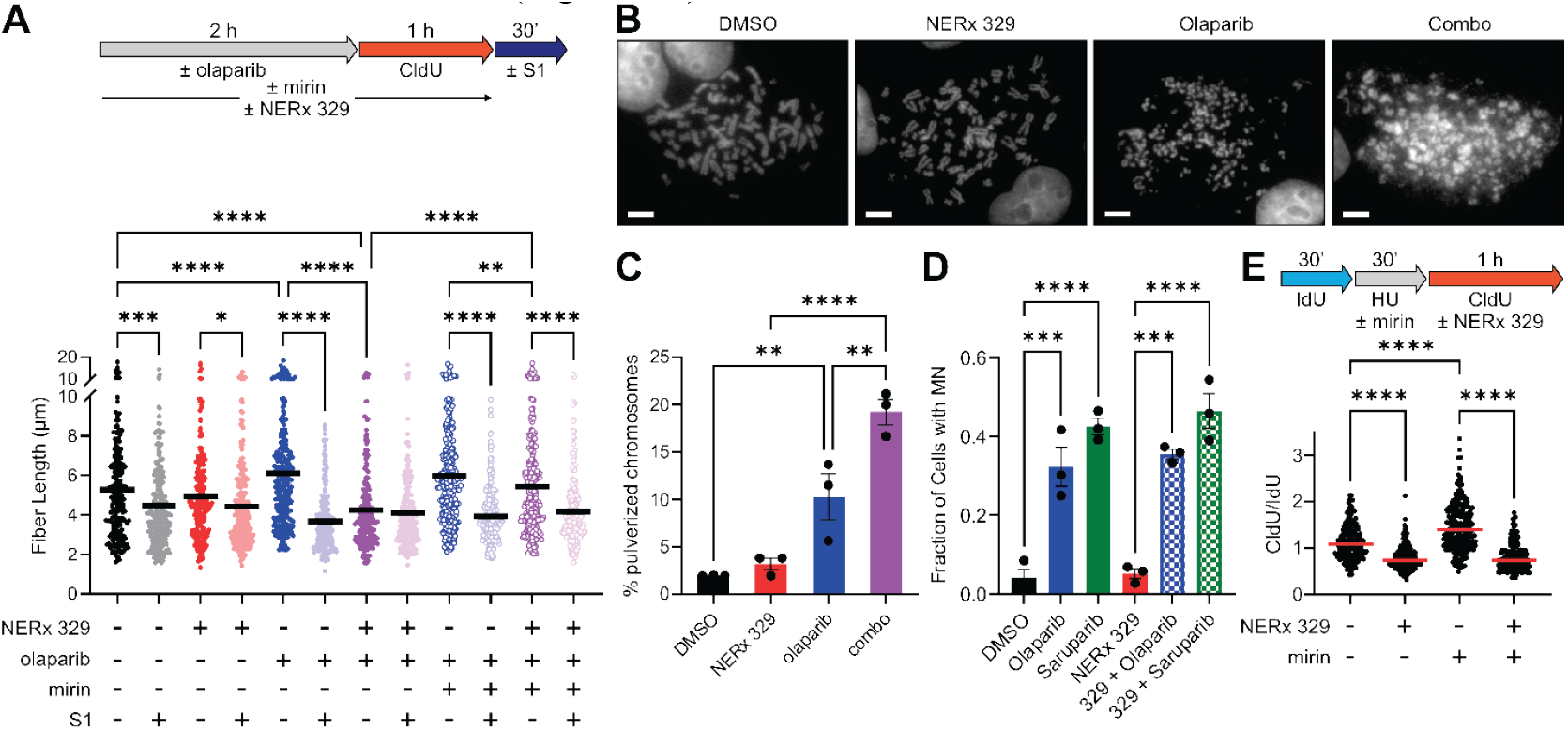
NERx-329 inhibits ssDNA gap protection through RPA exhaustion. (**A**) Labeling scheme (*top*) and quantification (*bottom*) of CldU fiber lengths from S1 nuclease DNA fiber combing experiments to measure replication fork progression and ssDNA gap formation. MDA-MB-436 cells were treated with vehicle control, NERx-329, olaparib, and/or mirin for 2 hr. before active replication forks were labeled with CldU for 1 hr. After processing, the purified DNA was incubated in the presence or absence of the S1 nuclease before the DNA fibers were combed and analyzed. The median measurement is indicated by the solid red bar (n = 205-250 fibers from duplicate experiments). (**B**) Representative metaphase spreads from MDA-MB-436 cells treated with vehicle control, NERx-329, olaparib, or their combination. (**C**) Quantification of pulverized chromosomes from the conditions depicted in panel B (n = 50 chromosomes; mean of triplicate experiments ± SEM). (**D**) Quantification of micronuclei formation in MDA-MB-436 cells treated with vehicle control or olaparib/saruparib in combination with NERx-329 (n = 100 cells, mean of triplicate experiments ± SEM). (**E**) Labeling scheme (*top*) and quantification (*bottom*) of DNA fiber combing experiments to measure replication fork restart after hydroxyurea treatment in MDA-MB-436 cells treated with vehicle control, NERx-329, and/or mirin. The median measurement is indicated by the solid red bar. All the data were analyzed via one-way ANOVA with Fisher’s LSD test, * P < 0.05, ** P < 0.01, *** P < 0.001, **** P < 0.0001.

Taken together, these data suggest that the amount of free RPA in BRCA1-deficient MDA-MB-436 cells is sufficient to protect the extensive amount of ssDNA gaps generated by olaparib treatment. However, combination treatment with NERx-329 exhausts RPA and inhibits ssDNA protection such that the abundant ssDNA gaps are degraded by MRE11. Notably, cells treated with RPAi alone still possessed ssDNA gaps (Figure 1A); therefore, the tested dosage and treatment duration did not completely exhaust RPA in the absence of additional replication stress-inducing agents. In contrast, olaparib treatment combined with NERx-329 yielded DNA fibers without gaps (Figure 1A), indicative of complete RPA exhaustion by the chemical inhibitor and unrestrained ssDNA gap formation. These data support the existence of an RPAi therapeutic window for cancer treatment afforded by oncogene- and/or DDRi-induced replication stress. Moreover, they describe a mechanism for the enhanced killing effect observed in NERx-329 and olaparib combination-treated BRCA1-deficient cancer cells. We hypothesize that enhanced killing is a result of genomic instability through replication fork collapse and chromothripsis, as has been observed under other treatment combinations that generate excessive ssDNA[19, 33, 34]. To evaluate this hypothesis, we performed metaphase spreads on single agent- and combination agent-treated MDA-MB-436 cells. Live-cell imaging experiments were performed to assess cell growth kinetics during single agent and combination treatments to determine the experimental timing for the detection of chromosome pulverization before significant levels of cell death were observed (Figure S1B). The results demonstrated that sequential treatment with olaparib for 2 days followed by NERx-329 for an additional day had a modest impact on growth on day 3 but completely inhibited growth by day 5. We employed this treatment scheme, and cells were collected for analysis of metaphase spreads on day 3 posttreatment (Figure 1B). Single-agent RPAi treatment had no observable effect on chromosome structure, whereas olaparib treatment increased chromosome pulverization. However, the combination strikingly induced chromosome pulverization (Figure 1B-C), presumably as a result of extensive replication fork collapse (Figure 1A). Taken together, NERx-329 chemically exhausts RPA such that olaparib-induced ssDNA gaps are degraded, and chromosomal integrity is compromised.

Considering the induction of chromosome pulverization, we assessed the generation of micronuclei (MN) with RPAi treatment in combination with olaparib and the PARP1-specific PARPi saruparib. MN in asynchronous cells was analyzed via DAPI staining, and the percentage of cells with at least 1 MN was quantified. The data revealed that PARPi treatment resulted in a significant increase in MN, whereas treatment with the single agent RPAi alone did not affect MN formation (Figure 1D). The combination of RPAi-PARPi treatment did not significantly alter the MN observed (Figure 1D). Taken together, these results indicate that MN formation is not required for the enhanced cell killing effect of the combination treatment.

Replication stress manifests in multiple ways that are measurable via the assessment of replication fork dynamics. The ability to restart stalled forks is essential for maintaining genomic stability. Therefore, we next evaluated the effect of RPA inhibition on replication fork restart. MDA-MB-436 cells were first grown with 5-iodo-2’-deoxyuridine (IdU) for 30 minutes, then replication was stalled with the addition of hydroxyurea (HU) for 30 minutes, and replication was resumed in the presence of CldU with and without NERx-329 for 1 hour (Figure 1E). As NERx-329 treatment results in the degradation of reversed forks[7] and BRCA-1 deficiency negatively impacts fork protection and restart[35, 36], both of which are dependent upon MRE11 nuclease activity, HU stalling was conducted in the presence and absence of mirin to ensure accurate measurement of replication fork restart as opposed to fork protection (Figure 1E). Similarly, mirin enhanced fork restart and increased the CldU/IdU ratio (Figure 1E). In both the presence and absence of mirin, treatment with RPAi yielded fibers with a decreased CldU/IdU ratio, indicative of impaired replication fork restart (Figure 1E). NERx-329-dependent inhibition of fork restart was independent of BRCA-1 status, as RPAi-treated NSCLC A549 cells also exhibited a decreased CldU/IdU ratio (Figure S1C). Collectively, these data reveal the impact of combination treatment focused on the disruption of replication fork dynamics and define a treatment scheme with potential therapeutic utility beyond BRCA-1 deficiency.

### *In vivo* anticancer activity of the RPA-PARPi combination therapy

To determine whether combination PARPi-RPAi treatment could be effective as a therapeutic strategy, we employed a BRCA1-deficient MDA-MB-436 TNBC cell line *in vitro* in combination with longitudinal live-cell imaging to assess cell growth. Olaparib did not impact proliferation until 48 hours after treatment, after which proliferation was reduced (Figure 2A), which is consistent with ssDNA gaps induced during S phase persisting through the cell cycle and forming DSBs in the second S phase[11]. RPAi-mediated inhibition of proliferation was evident within 24 hours, and cell confluence remained stable for ∼4 days, after which an increase was observed (Figure 2A). The efficacy of the single agent treatments was enhanced in combination, which resulted in essentially no growth observed through the 7-day assay and an actual reduction in confluence indicative of cell death and not simply a reduction in proliferation (Figure 2A). This implies that the PARPi-induced S phase ssDNA gaps that can persist through the next cell cycle are instead degraded in the initial S phase when RPA protection is concurrently inhibited, resulting in replication fork collapse and chromosome pulverization. The recovery of growth after ∼4 days of treatment with the single agent NERx-329 suggested either the emergence of resistance or a drug metabolism effect. To distinguish between these possibilities, we performed a similar experiment but added a second drug administration on day 3 (Figure 2B). The decrease in confluence upon a second NERx-329 dose demonstrated that the cells remained sensitive to RPAi treatment despite the initial resumption of cell growth (Figure 2B). These results and our previous *in vitro* analyses indicated that the combination of NERx-329 and olaparib could be an effective *in vivo* therapeutic strategy. To determine whether we could safely administer PARPi-RPAi treatment, we employed MDA-MB-436 models *in vivo* that were subcutaneously implanted into NSG mice. When the tumors reached ∼100 mm^3^, they were randomized to one of the 4 arms for single agent or combination treatment. Olaparib was administered once weekly, and RPAi was administered every 5 days on/two days off. We did not observe any increased toxicity in the combination-treated mice compared with the NERx-329-treated mice, as assessed by body weight measurements (Figure S2). The data presented in Figure 2C for assessing tumor volume over time show the efficacy of the combination, which recapitulates the live-cell imaging data where the combination abrogates tumor growth. The final tumor weight was also determined, and the efficacy of the combination treatment was confirmed (Figure 2D). These data demonstrate the utility of combination PARPi-RPAi therapy in BRCA1-deficient cancers.

**Fig. 2.**
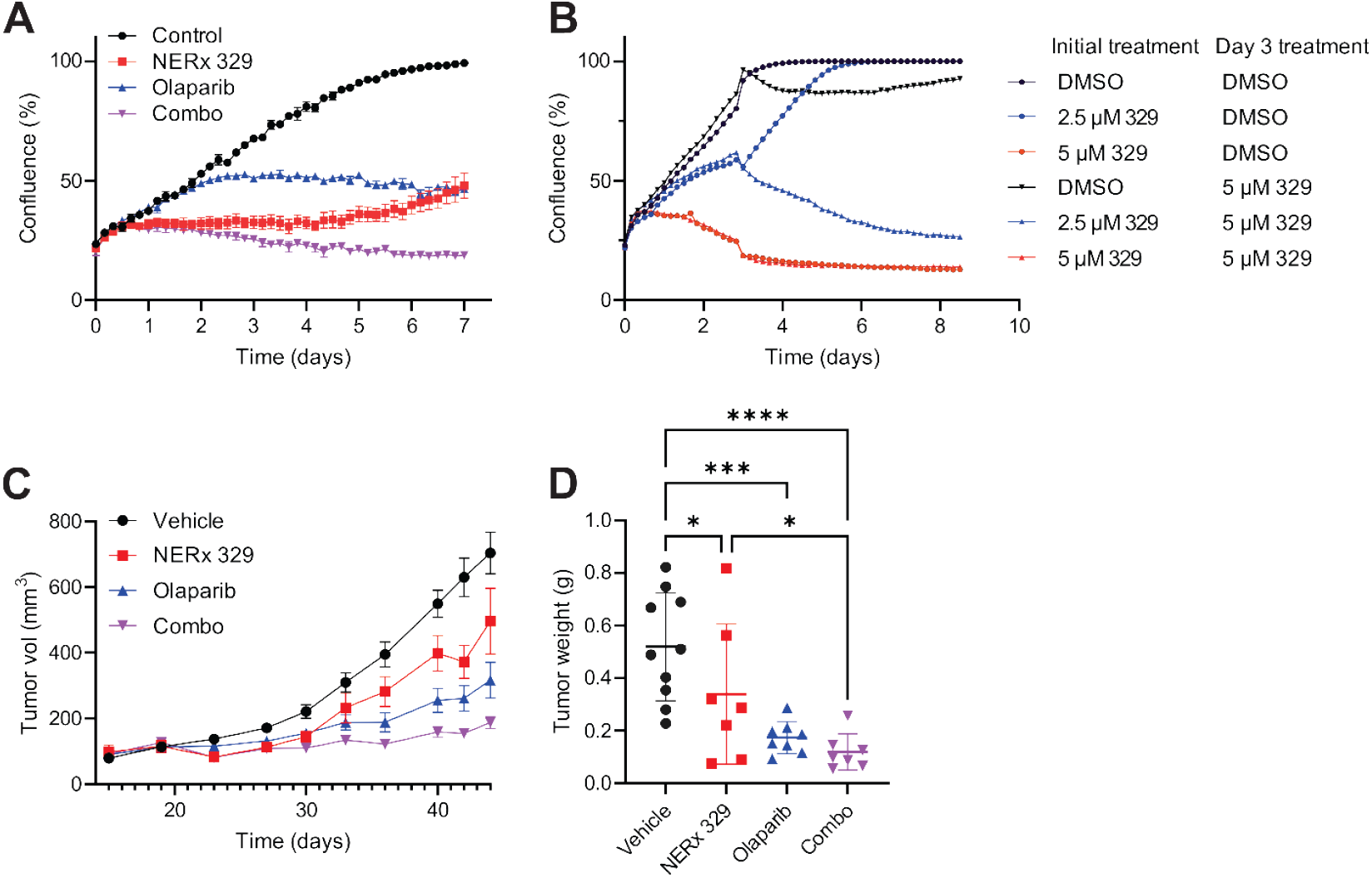
*In vivo* efficacy of the NERx329 and PARPi combination. (**A**) Confluence of MDA-MB-436 cells treated with vehicle control, NERx-329 (2 µM), olaparib (1 µM), or a combination of both as monitored by IncuCyte live-cell imaging. (**B**) Confluence of MDA-MB-436 cells treated on day 0 with vehicle control or NERx-329, followed by redosing treatment on day 3 with vehicle control or NERx-329 as monitored by Incucyte live-cell imaging. (**C**) Tumor volume growth over time in response to intraperitoneal injection of vehicle control, NERx-329, olaparib, or combination treatment in MDA-MB-436 cells subcutaneously implanted in NCG mice. (**D**) Terminal tumor weights from the experiment described in panel B. The average is indicated by the solid bar, and the data were analyzed by one-way ANOVA with Fisher’s LSD test; * P < 0.05, ** P < 0.01, *** P < 0.001, **** P < 0.0001.

### PARPi specificity and cancer genetics impact RPAi efficacy

As our mechanistic studies of ssDNA gap protection translated to a positive *in vivo* therapeutic combination with olaparib, we sought to determine how PARPi specificity impacted combination treatment. Olaparib is a potent inhibitor of both PARP1 and PARP2, while PARP1 is the most abundant PARP with known roles in gap suppression. To assess specificity, we employed the more potent and selective PARP1-specific inhibitor saruparib (AZD5305)[37]. Considering the accurate recapitulation of *in vivo* antitumor activity with 5-day live-cell imaging analyses, we assessed saruparib in combination with NERx-329 in the BRCA1-deficient MDA-MB-436 TNBC cell line via this methodology. The combination of RPAi treatment with the PARP1-specific inhibitor saruparib resulted in no growth and induced cell death, as indicated by Cytotox Red+ dye uptake (Figure 3A-B), similar to the results obtained with olaparib. Importantly, the increased potency of saruparib resulted in a 200-fold decrease in the concentration of the PARPi to 5 nM from the 1 µM needed for olaparib treatment of the same cells. These data demonstrate that the effects of chemical RPA inhibition are a result of PARP1 inhibition and ssDNA gap accumulation. To ensure that this was not a cell model-specific event and to determine the impact of BRCA1, we assessed the PARPi–RPAi interaction in the isogenic UWB1.289 ovarian cancer model[38]. In BRCA1-deficient UWB1.289 cells, the maximal activity of saruparib was observed at 50 nM (Figure 3C); a 10-fold increase to 500 nM had no additional effect on cell proliferation over the 7-day experiment (Figure S3A). NERx-329 significantly inhibited cell growth over the first 3 days of treatment (Figure 3C), after which proliferation resumed to a doubling time similar to that of the untreated cells, as previously observed (Figure 2A). In the combination treatment, the cells remained non-proliferative throughout the entire 7-day experiment despite a single treatment (day 0) (Figure 3C). These data support a model in which sufficient PARPi-dependent ssDNA is generated such that RPAi combination treatment induces chemical exhaustion of RPA and is responsible for the observed cell death. The BRCA1-complemented cell line is a model of PARPi resistance and, as expected, is far less sensitive to the PARPi saruparib; treatment at a dose of 25 µM had only a modest effect on cellular proliferation (Figure 3D). Importantly, these PARPi-resistant cells were still sensitive to RPAi single-agent treatment, and the combination clearly displayed greater anticancer efficacy than either individual treatment (Figure 3D). These data are consistent with PARPi induction of ssDNA gaps independent of BRCA1 status, with BRCA1-proficient cells requiring higher doses or longer PARPi exposure than BRCA1-deficient cells [9, 31]. Moreover, these data support the hypothesis that the combined efficacy of PARPi and RPAi is derived from processes that generate ssDNA and is not confined to the BRCA1-deficient background. To further evaluate this hypothesis, we used a CRISPR knockout screen to identify synthetic lethal interactors with NERx-329. The top hit was flap endonuclease 1 (FEN1), which is involved in Okazaki fragment maturation [39]. Consistent with the synergy of NERx-329 with genetic factors that impact ssDNA gap accumulation, FEN1 inhibition has been demonstrated to lead to ssDNA gap formation [27, 40]. To validate the FEN1-RPAi interaction, we used CRISPR/Cas9 gene editing to generate homozygous (FEN1 KO6) and heterozygous (FEN1 KO5) FEN1 knockouts in BRCA1 wild-type HR-proficient A549 NSCLC cells (Figure S3B). We confirmed that FEN1 KO cells were more sensitive to RPA inhibition than nontargeted control (NTC) cells were (Figure 3E-F) and that the NERx-329 response directly correlated with FEN1 expression levels (Figure 3E), owing to their intrinsic generation of lagging strand ssDNA gaps. These results highlight the key role that RPAi can play when certain factors that influence ssDNA gap formation are inactivated (BRCA1, PARP1, FEN1) owing to genetic background or drug exposure. The NERx-329 therapeutic window is derived from oncogenic replication stress; thus, other processes, either genetically driven or exogenously driven, that generate ssDNA gaps further increase RPAi potency.

**Fig. 3.**
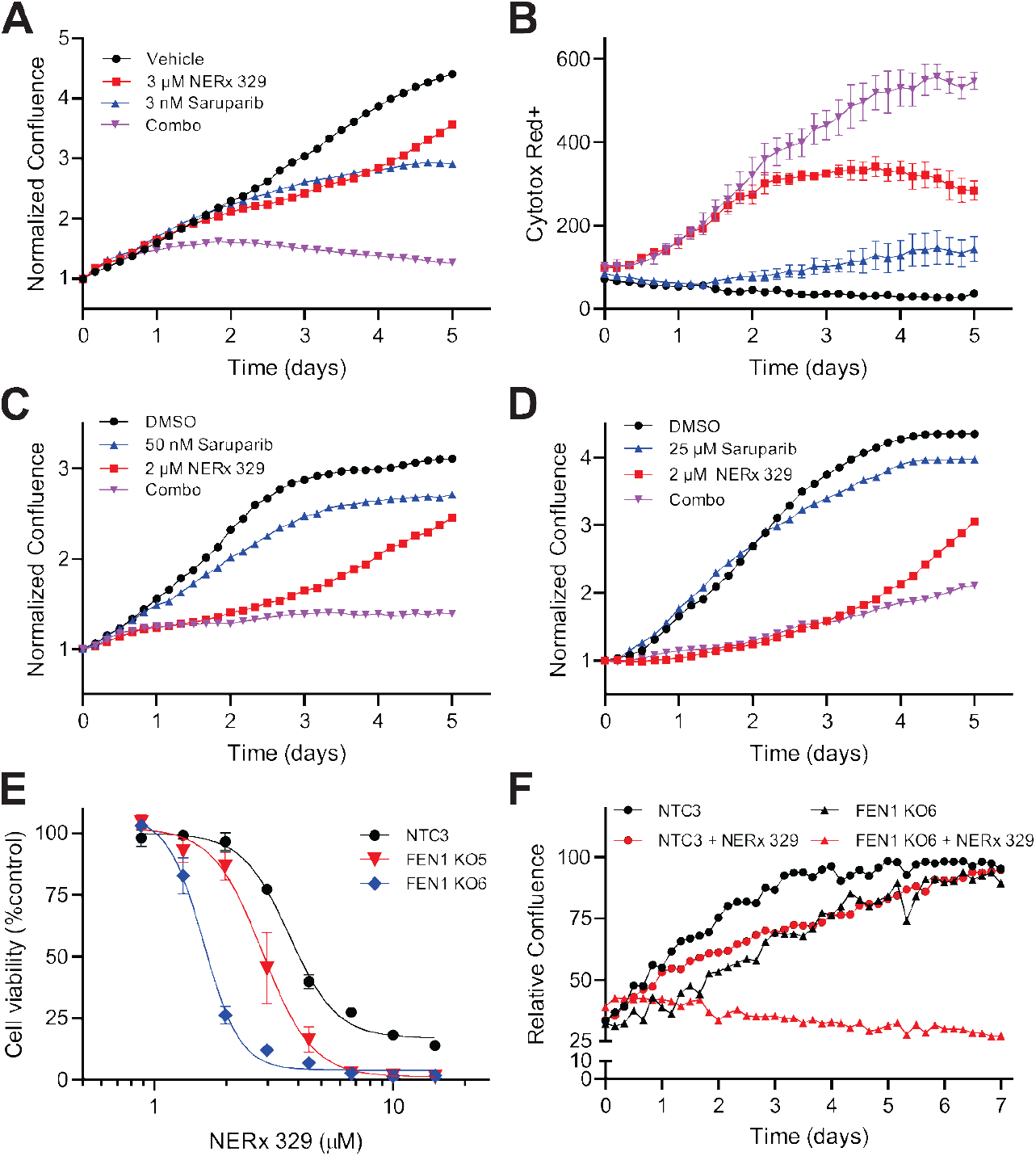
NERx-329 potency upon loss of ssDNA gap-suppressing factors. (**A**) Confluence of MDA-MB-436 cells treated with vehicle control, NERx-329, saruparib, or their combination as monitored by IncuCyte live-cell imaging. (**B**) Cytotox Red+ dye fluorescence intensity from the experiment depicted in panel A. (**C**) Confluence of UWB1.289 cells treated with the vehicle control, NERx-329, saruparib, or combination as monitored by IncuCyte live-cell imaging. (**D**) Confluence of BRCA1-complemented UWB1.289 cells treated with vehicle control, NERx-329, saruparib, or combination as monitored by Incucyte live-cell imaging. (**E**) Viability of A549 NTC3, homozygous FEN1 KO6, and heterozygous FEN1 KO5 cells treated with NERx-329 as assessed by a CCK-8 assay. (**F**) Confluence of A549 NTC3 and FEN1 KO6 cells treated with vehicle control or NERx-329 as monitored by IncuCyte live-cell imaging.

### Impact of RPAi-PARPi combination therapy on cell cycle progression

To better understand the downstream cellular effects of the loss of ssDNA gap protection that underlies the efficacy of the RPAi combination *in vitro* and *in vivo*, we investigated its impact on cell cycle progression. Previous analyses of predecessor RPAi demonstrated minimal effects on the cell cycle in BRCA wild-type cancers [21], and the results obtained in BRCA1-deficient MDA-MB-436 cells were very similar, with minimal impact observed (Figure 4A). PARPi treatment in BRCA1-proficient cell lines results in the accumulation of cells in G2/M with a reduction in the G1 population[31], whereas BRCA1-deficient ovarian cancer cells do not due to a failure in checkpoint activation [11]. Similarly, we found that olaparib single-agent activity slightly increased the number of cells in the G2/M phase and decreased the number of cells in the G1 phase (Figure 4A). All the observed changes in the cell cycle distribution were driven by PARPi, as combination treatment with RPAi and PARPi did not further affect the cell cycle distribution (Figure 4A-B). To assess the effects on individual cells more accurately, we employed the fluorescent ubiquitin cell cycle imaging (FUCCI) system [41] to monitor cell cycle progression in live MDA-MB-436 cells (Supplemental Movies 1 and 2, DMSO- and RPAi-treated, respectively). FUCCI cells exhibit red fluorescence during G1 (Cdt1-mKate2) and green fluorescence during S-G2-M (geminin-mTagGFP2) [42, 43]. The cells thus are fluorescent yellow at G1/S and are colorless during division and initiate the next cell cycle (M/G1). The growth of MDA-MB-436 FUCCI cells in response to RPAi, saruparib, or the combination treatment was similar to that of the parental MDA-MB-436 cell line (Figure S4A). We found that NERx-329 had a biphasic effect on the distribution of the MDA-MB-436 FUCCI cell cycle (Figure 4B-D). The initial phase exhibited a modest increase in the S-G2-M population at the expense of the G1/S population over ∼48 hours (Figure 4C-D). The second phase exhibited a large accumulation of the G1 population (Figure 4B) concomitant with a significant reduction in the S-G2-M population (Figure 4D). Both single agent treatment and combination treatment with saruparib had little to no effect, and disruption of the cell cycle was driven by NERx-329 treatment in a dose-dependent manner (Figure 4B-D, S4B-E). Similar G1 accumulation was observed in NSCLC A549 FUCCI cells (Figure S4F-H), indicating that the effects of the cell cycle were independent of BRCA1 status. To further evaluate these RPAi-dependent effects, we tracked individual vehicle control- or NERx-329-treated MDA-MB-436 FUCCI cells for 72 hours or until they could no longer be tracked due to cell overlap (Figure 4E). We found that NERx-329 treatment induced G1 and S-G2-M delays in a dose-dependent manner, with a minor effect on M/G1 progression (Figure 4E-F). Consistent with the induction of replication catastrophe, the prolonged G1 effect was dependent on RPAi treatment occurring in the previous S phase; MDA-MB-436 FUCCI cells that were in M/G1 phase at the initiation of RPAi treatment did not exhibit delayed G1 durations compared with those starting in G1, G1/S, or S-G2-M, all of which would be required to go through S phase before presenting a measurable G1 phase (Figure S4I). We also noticed that a substantial portion of the MDA-MB-436 FUCCI cells treated with NERx-329 exhibited mitotic bypass by transitioning from green to colorless to red throughout the next cell cycle without successfully dividing (Figure 4G, Supplemental Movie 3). We did not observe evidence of abortive mitosis or binuclear formation in the G1 population of these cells but rather an increased area of nuclear Cdt1-mKate2 expression after 72 hours of treatment (Figure S4J-M). Analyzing these results within the context of the single-cell trajectories revealed that cells that were in the later stages of S-G2-M at the start of RPAi treatment (those that underwent mitosis in <10 hrs.) were able to successfully divide, whereas those that were in the earlier stages of S-G2-M at the start of RPAi treatment (those that underwent mitosis in >10 hrs.) exhibited a drastically increased frequency of mitotic bypass (Figure 4E). These results are consistent with other reports describing the S phase- and DDR-dependent induction of mitotic bypass in response to oncogenic replication stress or other DNA-damaging agents [44-47].

**Fig. 4.**
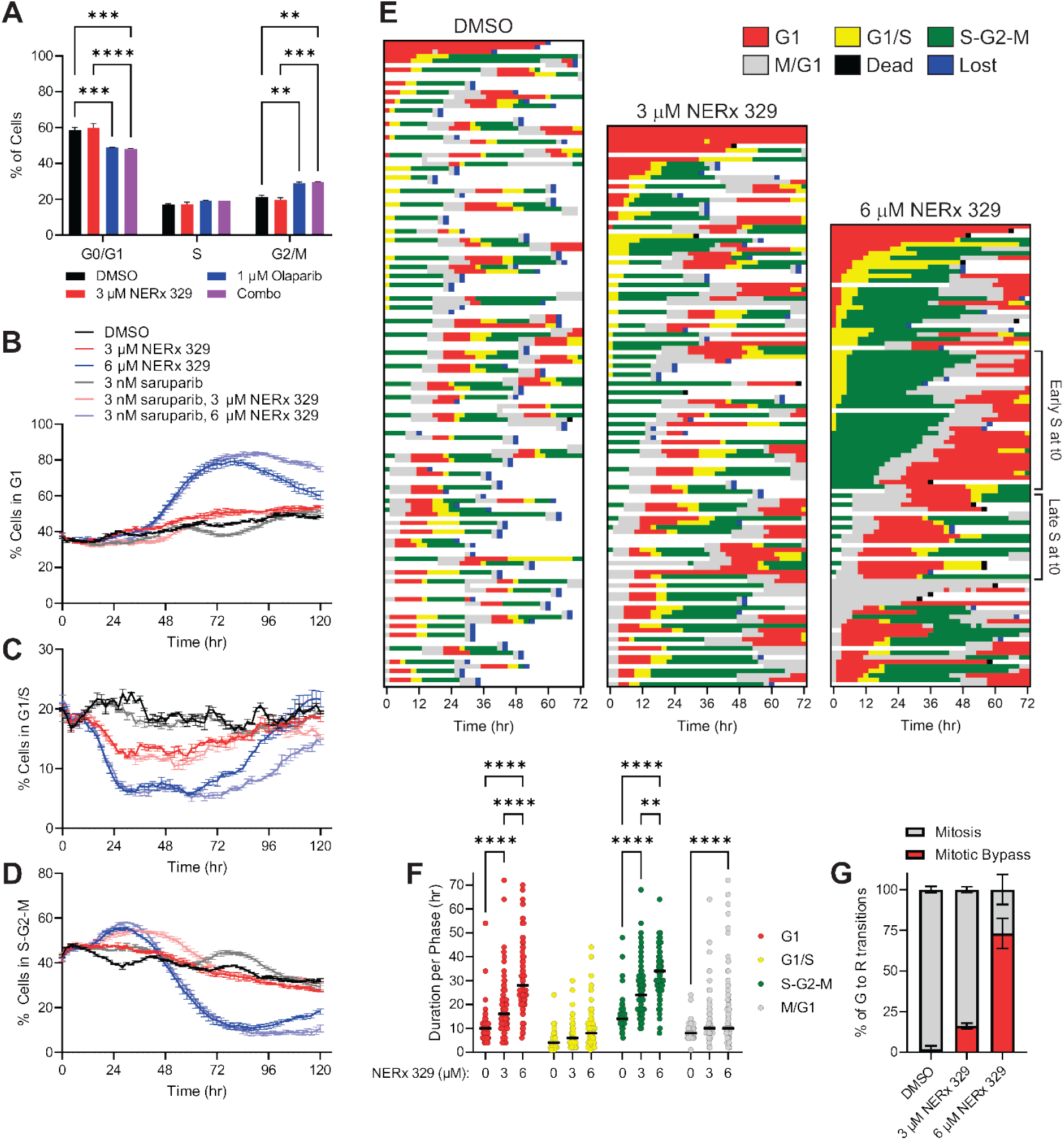
Effects of NERx-329 and PARPi combinatin treatment on the cell cycle. (**A**) Quantification of the cell cycle distribution of MDA-MB-436 cells treated with vehicle control, NERx-329, olaparib, or their combination, as measured by flow cytometry (mean ± SEM). Quantification of MDA-MB-436 FUCCI cells in (**B**) G1, (**C**) G1/S, or (**D**) S-G2-M following treatment with vehicle control, NERx-329, saruparib, or the combination (mean ± SEM). (**E**) Individual MDA-MB-436 FUCCI cell trajectories following treatment with vehicle control or increasing concentrations of NERx-329 (n = 54, 68, and 85 cells at the initial time points, respectively, from duplicate experiments). Single cells were followed until they were lost/untrackable due to cell overlap (blue). (**F**) Quantification of the MDA-MB-436 FUCCI cell cycle duration following treatment with vehicle control or increasing concentrations of NERx-329 (n = 66-112 from duplicate experiments). The median measurement is indicated by the solid black bar. (**G**) Quantification of MDA-MB-436 FUCCI cells that transitioned from green (G) to red (R) and successfully divided (mitosis) or engaged in the next cell cycle without dividing (mitotic bypass) following treatment with vehicle control or increasing concentrations of NERx-329 (n = 34-50 transitions from duplicate experiments). The data were analyzed via one-way ANOVA with Fisher’s LSD test, * P < 0.05, ** P < 0.01, *** P < 0.001, **** P < 0.0001.

### RPAi and PARPi as predictive tools for cancer patient treatment

The previous results describe the *in vitro* use of pharmacological tools within certain genetic backgrounds as predictors for *in vivo* responses that could also be translated to patient care. To assess clinically relevant models, we established a panel of twelve cell lines from high-grade serous ovarian cancer (HGSOC) patients. Ascites fluid was collected, and the cell lines were established as adherent monolayers that were CK7- and EpCAM-positive, indicative of HGSOC (Figure S3A). Despite advances in and the advent of PARPi, cisplatin remains the backbone treatment for most HGSOC patients [48]. We therefore first evaluated patient-derived cell line sensitivity to cisplatin in spheroids and then quantitative sensitivity to cisplatin and RPAi in monolayer cultures. The individual patient-derived cell lines displayed sensitivity to both agents (Figure S3B-C) over a fairly narrow range, with HDH5 exhibiting a sensitive phenotype. Whole-genome and whole-exome sequence analyses revealed a series of mutations in DNA damage and repair genes that could impact RPAi sensitivity and PARPi-RPAi combination efficacy, although no known deleterious BRCA1 mutations were identified (Figure S4D). The HDH5 cell line contains a benign BRCA1 somatic mutation, G890V [National Center for Biotechnology Information. ClinVar; VCV000037479.55], and a potentially deleterious BRCA2 splice site mutation, 632 G-A [National Center for Biotechnology Information. ClinVar; VCV000052065.12] (Figure S4D, Table S1). Characterization of this cell line as spheroids revealed sensitivity to NERx-329 or cisplatin, with IC_50_ values slightly higher than those observed for analysis in monolayer cultures (Figure 5A-B). Imaging analyses revealed that the size of the spheroids did not appreciably change over the 7-day time course in vehicle-treated cells, but the spheroids became denser as the cells divided (Supplemental Movie 4). This is consistent with the recent description of tumor structure involving cytocapsular structures that can restrict size[49]. Treatment with NERx-329 induced a decrease in cell viability, as assessed by Cytotox Red+ dye uptake, which is indicative of the loss of membrane integrity (Figure 5C, Supplemental Movie 5). Interestingly, the size of the spheroids treated with NERx-329 increased with increasing red intensity both over time and as a function of the NERx-329 dose (Figure 5C), which was again consistent with cell death, loss of membrane integrity and dissolution of the restrictive cytocapsular structures [49]. Loss of the cytocapsular structure and loss of cell viability in HDH5 cells and other patient-derived spheroid models (HDHs 9 and 11) were also observed in response to cisplatin treatment (Figure S4a).

**Fig. 5.**
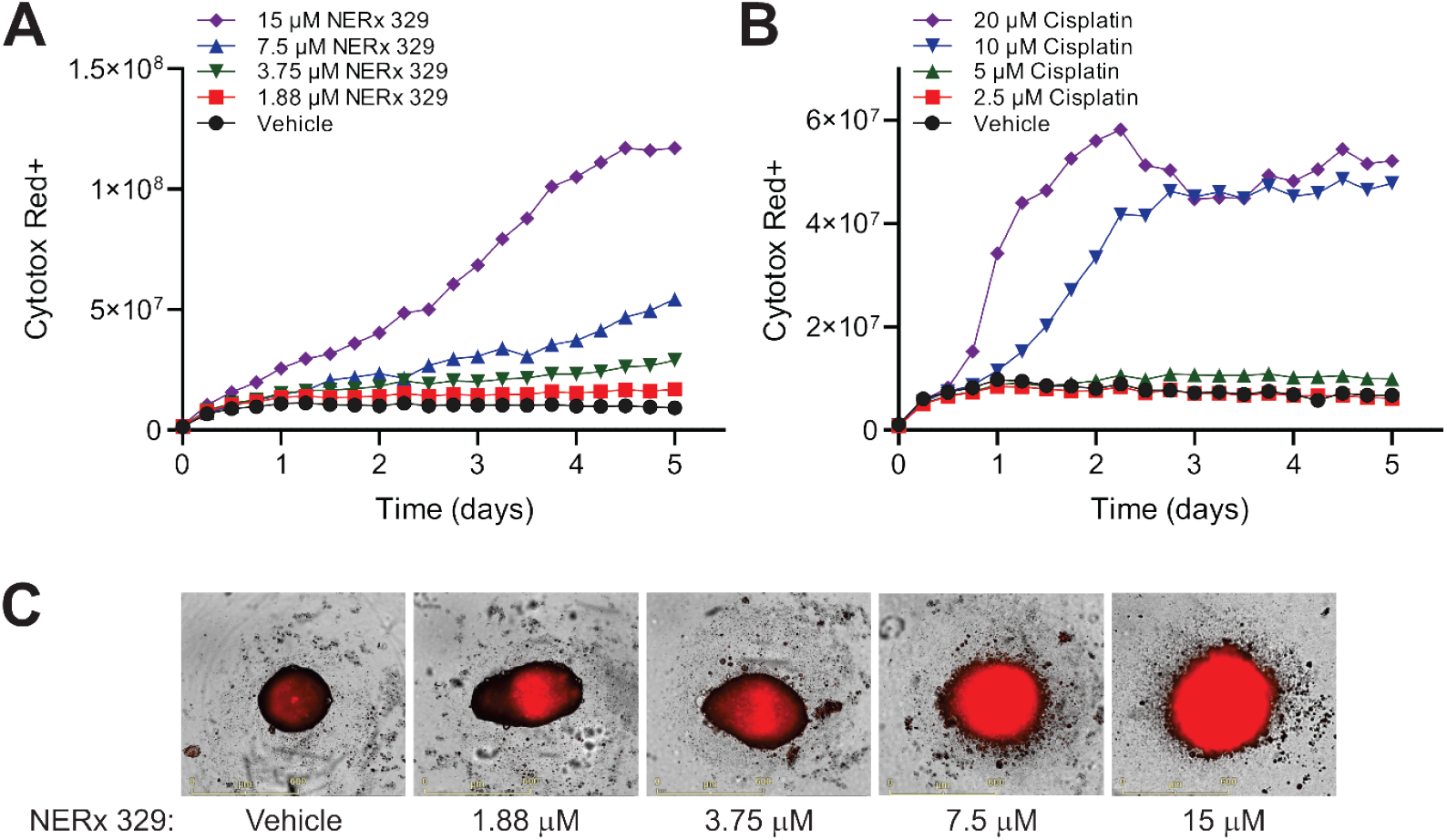
Sensitivity of HGSCO HDH5 spheroids to NERx-329 and cisplatin. (**A**) Cytotox Red+ dye fluorescence intensity of HDH5 spheroids treated with increasing concentrations of NERx-329 as monitored by Incucyte live-cell imaging. (**B**) Cytotox Red+ dye fluorescence intensity of HDH5 spheroids treated with increasing concentrations of cisplatin as monitored by Incucyte live-cell imaging. (**C**) Representative Cytotox Red+ images of HDH5 spheroids treated with NERx-329.

The BRCA2 mutation in HDH5 cells has been characterized as potentially deleterious; therefore, we assessed sensitivity to the PARP1i saruparib alone and in combination with RPAi. HDH5 cells are relatively resistant to saruparib, with no significant decrease in cell growth at the relatively high concentration of 5 µM (Figure 6); in comparison, the proliferation of BRCA1-deficient MDA-MB-436 and UWB1.289 cells was altered by single-digit nanomolar saruparib treatment (Figure 2A, 3A, respectively). These data suggest that BRCA2 mutation does not alter ssDNA gap suppression, replication fork protection, or HR-dependent DSB repair, all of which have been implicated in driving sensitivity to PARP inhibition[50, 51] and are therefore unlikely to be deleterious. Interestingly, HDH5 cells are more sensitive to NERx-329 treatment (Figure 6), whereas both the MDA-MB-436 and UWB1.289 cell lines showed a significant but modest effect on cell growth when treated with 2 and 3 µM NERx-329, respectively. HDH5 cells are extremely sensitive to NERx-329 treatment at 2 µM, which dramatically affects cell growth and death. In addition, HDH5 cells were extremely sensitive to the combination of PARPi and RPAi treatment. These data suggest that the BRCA2 632 G-A splice site mutation may impact RPAi single-agent and PARPi combination efficacy but does not affect single-agent PARPi activity. However, HDH5 also has mutations in other DDR genes, including replicative DNA polymerases δ and ε, which could account for the sensitivity of HDH5 to the combination of RPAi and PARPi-RPAi (Figure S4D, Table S1). However, the lack of single agent PARPi sensitivity in HGSOC patients harboring the BRCA2 632 G-A mutation suggests that PARPi are not part of the treatment regimen for HGSOC patients harboring this specific BRCA2 mutation. These data also suggest the utility of RPAi treatment for cancers that are/have become resistant to PARPi and illustrate a more precise way to identify and effectively treat patients who may not respond to typical PARPi treatment.

**Fig. 6.**
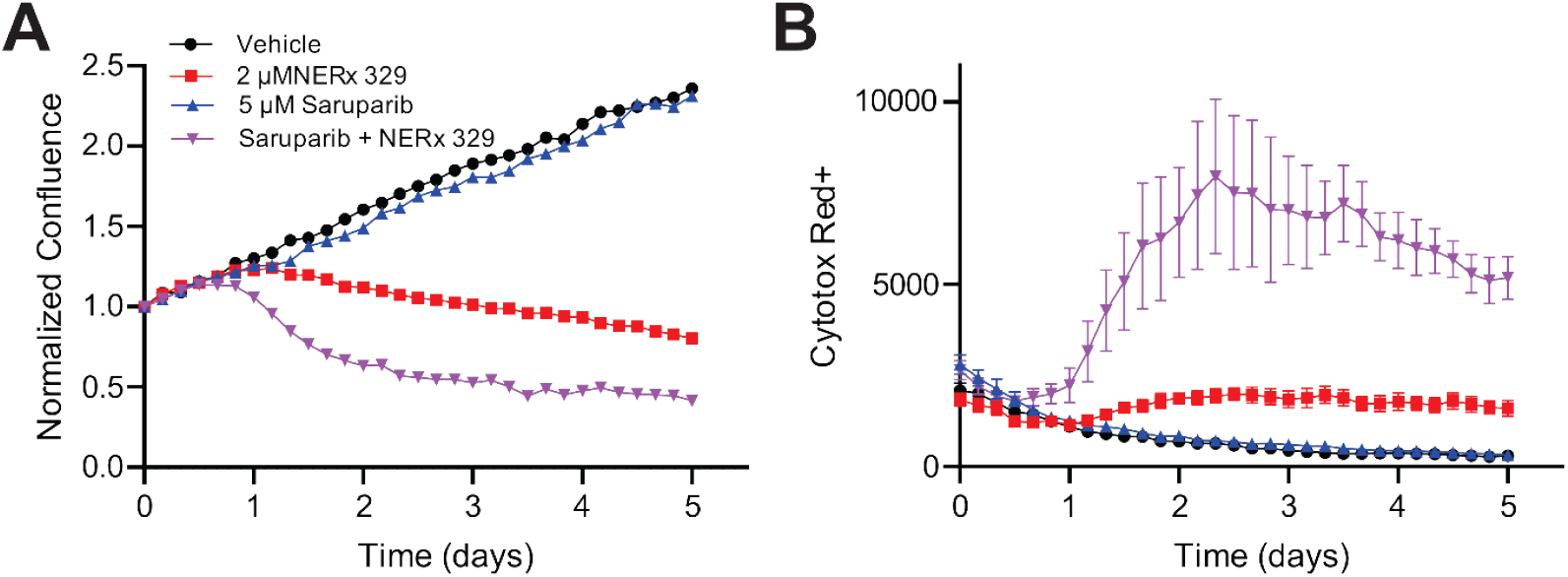
Sensitivity of HGSOC HDH5 cells to NERx-329-PARPi combination treatment. (**A**) Confluence of HDH5 monolayer cells treated with vehicle control, NERx-329, saruparib, or their combination as monitored by Incucyte live-cell imaging. (**B**) Cytotox Red+ dye fluorescence intensity from the experiment depicted in panel A.

## DISCUSSION

Nearly two decades ago, synthetic lethality was first demonstrated for the use of PARPi in BRCA-deficient cancers [52, 53]. Cell death is attributed to the formation of DSBs via replication through unrepaired single-strand nicks in the template DNA that cannot be repaired HR due to the deficiency of BRCA proteins. The discovery of PARP trapping activity as a driver of cellular sensitivity added the additional caveat that PARPi may induce a type of DNA damage through the direct trapping of PARP on DNA [54]. While the importance of BRCA protein-dependent processes is certainly an aspect of PARPi lethality [50], recent investigations have uncoupled PARPi sensitivity from HR/DSB repair and instead pointed to ssDNA gap accumulation as the initial driver of the therapeutic response [9, 30, 55]. In this model, ssDNA gaps are proposed as the key therapeutic lesion-driving sensitivity, whereas DSBs are in part downstream manifestations resulting from apoptosis [9, 11]. While the absolute size of ssDNA gaps that are formed in cancer cells is not known and likely differs depending on drug combination and genetic background, ssDNA gaps have been observed to be between 100–2000 basses when induced in eukaryotic systems [56-58]. Moreover, the combination of multiple confounding factors (in our case, BRCA1 deficiency and PARPi) increased the frequency of ssDNA gap occurrence and size [57]. Taken to extremes, these compounding effects can induce sufficient ssDNA gaps to sequester and exhaust all RPA such that unprotected ssDNA leads to replication fork collapse and chromothripsis [19]; alternatively, the same result can be achieved via RPAi combination treatment (Figure 7).

In this work, we delineated the underlying *in vivo* mechanism of the RPAi-PARPi combination in BRCA1-deficient cancer as the loss of ssDNA gap protection. These studies demonstrate that the efficacy of PARP-targeted therapy is enhanced by depleting RPA and lowering the ssDNA protection threshold (Figure 7). Therefore, both treatments individually function to target the protection of ssDNA; PARPi induce ssDNA and RPAi deplete the RPA protection capacity. When combined, the mechanism of the synergistic interaction involves contributions from both BRCA1 deficiency and PARPi treatment, which generate excess ssDNA through the formation of lagging strand gaps [14], and RPAi treatment, which sequesters free RPA, leading to RPA exhaustion (Figure 7). In this context, RPAi treatment exploits specific vulnerabilities for anticancer action and offers the potential to reduce PARPi dosage to limit PARPi-associated off-target toxicity or to overcome PARPi resistance, which is almost universally observed in patients with BRCA1-mutant HGSOC [59]. Moreover, the increased sensitivity to RPAi via ssDNA gap induction in three distinct backgrounds, deficiency of BRCA1 or FEN1 and inhibition of PARP1, suggests that other genetic alterations that impact RF dynamics and induce ssDNA gaps may also be susceptible to RPAi (Table S2). RPA inhibition functions through the same mechanism as described in this work to sequester free RPA alongside the ssDNA generated by therapeutic/deficiency (Figure 7). This may therefore allow for lower, more tolerable drug doses by minimizing off-target toxicities when used in combination with RPAi. Additionally, the identification of genetic factors that drive ssDNA gap formation could lead to novel therapeutic targets for use in combination studies. Moreover, potential NERx-329 therapeutic combinations provide novel alternatives to systems of acquired resistance, e.g., PARPi resistance, as RPAi combination efficacy is not dependent upon PARP function specifically but rather on ssDNA formation and protection in general. Indeed, PARPi-resistant cells exhibit ssDNA generation due to the expansion of nicks/SSBs, which can likely be leveraged for RPAi therapeutic efficacy [12]. RPA exhaustion-driven cancer cell death therefore could be useful in many cancers that arise from genomic instability initiated by mutations or alterations in genes that participate in maintaining genomic integrity.

## MATERIALS AND METHODS

### Cell Culture

MDA-MB-436 and MD-MB-468 cells were cultured in 1:1 DMEM: Ham’s F12 (Corning) supplemented with 10% FBS and penicillin/streptomycin. UWB1.298 cells were grown in 1:1 MEGM (Lonza): RPMI (Corning) supplemented with 3% FBS. For UWB1.289 BRCA1 complemented cells, 200 µg/mL G-418 (Sigma) was added to the media for maintenance cultures but not when the cells were treated. HDH5 cells were grown in modified NOSE-CM [24], 1:1 Media 199:MCDB 105 (Sigma-Aldrich) with 2.2 g/L NaHCO_3_, 10% FBS, and components of the MEGM Mammary Epithelial Cell Growth Medium SingleQuots Kit (Lonza, Catalog #: CC-4136), with 1 kit/liter of media. All the cell lines were grown in a CO_2_ incubator at 37°C.

### DNA Fiber Combing

In brief, MDA-MB-436 or A549 cells were treated as indicated to label active replication tracks with IdU and/or CldU, collected, and lysed by proteinase K digestion in agarose plugs. DNA was released by melting the agarose plugs and fibers were combed onto coverslips with a FiberComb machine (Genomic Vision). Fibers were immunostained with antibodies towards IdU or CldU and fiber length was measured using ImageJ. For S1 fiber combing assays, collected cells from each condition were split into two agarose plugs and processed identically until DNA was released, at which point one sample was treated with S1 nuclease and the other was mock treated before being combed onto coverslips. See Supplemental Material and Methods for full experimental details.

### Metaphase Spreads

For metaphase spread experiments, 2 × 10^5^ MDA-MB-436 cells were plated in a 6-well plate and grown for 24 hr. at 37°C with 5% CO_2_. After 24 h, the cells were treated with 2 µM olaparib or the DMSO control for 2 days at 37°C with 5% CO_2_. Either 5 or 10 µM NERx-329 or the DMSO control was added for 1 additional day at 37°C with 5% CO_2_. The cells were then treated with colcemid for 3 h at 37°C with 5% CO_2_. The media was removed, the cells were washed twice with PBS, and the cells were trypsinized. All media and washes were combined with the trypsinized cells and centrifuged. The cells were then incubated in 0.56% KCl (w/v) for 15 min at room temperature. The cell suspension was centrifuged, and the cells were fixed in methanol: glacial acetic acid (3:1) on ice for 1 hr. The fixed cells were centrifuged and resuspended in a small volume of fixative solution and then dropped onto alcohol-cleaned slides from a height of 6–12 inches. The slides were allowed to dry at room temperature for 30 min and then at 42°C for 10 min. Coverslips were mounted using Vectashield antifade mounting medium with DAPI. Chromosomes were visualized using an EVOS FL Auto 2 Imaging System (Invitrogen) under a 60X oil immersion lens using a DAPI filter. For each replicate experiment, 50 chromosomes were counted for each experimental condition.

### Micronuclei quantification

MDA-MB-436 cells were plated in 8-well chamber slides (CellTreat #229168) at 4 × 10^4^ cells per chamber and allowed to adhere overnight. The cells were treated with the RPAi NERx-329, the indicated PARPi, and drug combinations for 2 hours as indicated. The amount of DMSO used was held constant at 0.5% for all the treatments. After the 48-hour treatment, 3.5 µg/mL cytochalasin-B in 0.5% DMSO was added to each well to arrest the cell cycle. After 24 hours of cytochalasin-B exposure, the media and chambers were removed, and the cells were incubated with a solution of 45% PBS, 45% 0.075 µM KCl, and 10% 25:1 methanol:acetic acid for 10 minutes, followed by another 10-minute incubation with only the 25:1 methanol:glacial acetic acid mixture. The slide was then washed 3 times with PBS, and a coverslip was mounted using Vectashield H-1500 with DAPI and allowed to cure at room temperature for 15 minutes before being stored overnight at 4°C in the dark. Images were captured with an EVOS FL Auto2 microscope, and an average of 100 cells were scored manually for the presence of micronuclei, following established parameters [25].

### Live cell analysis

For adherent live cell analysis, cells were plated at 5×10^3^ cells/well in a 96-well plate and incubated for 18–24 hours in a CO_2_ incubator at 37°C. Treatments were as indicated in the figures. If indicated, IncuCyte® CytotoxRed Dye (Sartorius, Catalog#: 4632) was added at a final concentration of 0.25 µM at the time of drug treatment. The cells were transferred to the IncuCyte imager after treatment, and images were captured at 4X or 10X magnification at the time intervals indicated. Confluence was determined via IncuCyte software analysis. Cytotox Red dye uptake was measured by the total integrated red fluorescence intensity. To assess spheroid culture viability, 1×10^4^ cells/well were plated in ultralow attachment 96-well plates (Corning, Catalog#: 7007) and incubated for 24–48 hours until spheroids formed. The cultures were then treated as indicated and transferred to an IncuCyte Imager. Images were captured at 4X or 10X magnification at the indicated time intervals, and CytotoxRed uptake was measured as the total integrated intensity of the red fluorescence within the spheroid boundary.

### In vivo analysis

Female NOD-scid/IL2Rg-null (NSG) mice (*In Vivo* Therapeutics Core Facility, IU Simon Comprehensive Cancer Center, Indianapolis, IN, USA) were used and housed in a pathogen-free facility at the IUSM LARC facility. NSG studies were approved by the Institutional Animal Care and Use Committee at Indiana University School of Medicine. The hind flanks of 8–10-week-old NSG mice were implanted with MDA-MB-436 cells (∼2.5 × 10^6^) in Matrigel. The tumor volume was monitored via electronic calipers [tumor volume = length × (perpendicular width)^2^ × 0.5]. Mice with tumors ranging from ∼100 mm^3^ in size were randomized into individual treatment arms. RPAi NERx-329 was formulated in 5% NMP, 60% PEG300, and 30% Tween-80 and administered via intraperitoneal (IP) injection at 25 mg/kg once daily. Olaparib was dissolved in PBS and administered by oral gavage at 5 mg/kg once daily. Tumor volumes were monitored as indicated, and the results are presented as the average tumor volume ± standard error of the mean for each group. Tumor weight was determined at the end of the experiment from excised tumors. The number (n) for each experiment is presented in the figure legend.

### Ovarian Cancer PD Spheroid Models

Primary EOC cells from ascites were maintained in NOSE-CM [24] with minor modifications as described in the Cell Culture section above. Ultralow attachment (ULA) plates were obtained from Corning. Patient samples were collected at Indiana University Hospital via protocols approved by the Indiana University Institutional Review Board (IRB). Excess ascites not required for clinical pathologic analysis were collected from study participants already undergoing cytoreductive surgery for suspected EOC. The ascites was transported to the laboratory at room temperature and processed to establish short-term 2D cultures of EOC as described by Shepherd et al. [24] or direct to spheroid (DTS) cultures. For DTS processing, ascites was mixed with RBC lysis solution, rocked for 30 minutes, and centrifuged at 200 × g for 10 minutes, after which the cell pellet was suspended. This process was repeated cell pellet was suspended in modified NOSE-CM for plating directly into spheroids on ULA plates. The cells were also frozen at this point and could be resurrected for growth in monolayers or spheroids as needed. Following the establishment of the cultures, primary EOC cells from ascites were maintained in NOSE-CM culture media [26] and subcultured at approximately 90% confluence. Upon passaging, the cells were split and plated at a 1:2 to 1:3 ratio, and aliquots were frozen in liquid N2. Short-term expanded cultures were utilized to establish 2D and 3D cultures and for analysis of drug sensitivity. Short-term cultures were routinely conducted for 3–4 passages and discarded if 90% confluence was not reached within 96 hours from a 1:2 or 1:3 dilution.

## List of Supplementary Materials

Supplemental Materials and Methods

Fig. S1 to S5

Data file S1

Movies S1 to S5

References (*1*)

## Acknowledgments

The authors thank Dr. Lata Balakrishnan and the members of the Turchi Lab for critical reading of the manuscript and the IUSCCC Preclinical Modeling and Therapeutic Core *in* Vivo Team.

## Author contributions

Conceptualization: PSV-C, MRJ, JEG, HDH, JJT

Methodology: PSV-C, MRJ, JEG, HDH, XS, SL

Investigation: PSV-C, KEP, JW, KSP, JTT

Visualization: PSV-C, MRJ, JEG, HDH, XS, SL

Funding acquisition: KSP, JTT

Project administration: JJT

Supervision: JJT

Writing – original draft: MRJ, JTT

Writing – review & editing: All authors

## Data and materials availability

All data are available in the main text or the supplementary materials.

