## Supplemental data for "Chemical inhibition of RPA gap protection sensitizes BRCA1-deficient cancers to PARP inhibition"

### Supplemental Materials and Methods

#### *DNA Fiber Combing*

For the S1 nuclease fiber combing experiments, 5 × $10^5$ MDA-MB-436 cells were plated in a 6-well plate and grown for 24 hr at 37°C with 5% $CO_2$. The cells were then treated as indicated with DMSO, 10 µM olaparib, 50 µM mirin, and/or 30 µM NERx 329 for 2 hr. The final concentration of DMSO was 0.7% across all treatment combinations. After 2 hr, the media was aspirated, and fresh media was added as indicated with DMSO, 250 µM CldU, 50 µM mirin, and/or 30 µM NERx 329 for 1 hr. The cells were then washed three times with prewarmed PBS and trypsinized. The cells from each condition were then evenly split into two agarose plugs (for eventual ± S1 nuclease treatment) and lysed overnight with 2 mg/mL proteinase K (Thermo Fisher) at 50°C. The next day, the agarose plugs were washed three times with TE buffer for 1 h each at room temperature with gentle rotation. Agarose plugs were then added to 50 mM MES (pH 5.5) and 100 mM NaCl at 68°C for 20 min, followed by incubation at 42°C for 10 min. β-agarase (New England Biolabs) was added to digest all the agarose and release the DNA overnight at 42°C. The next day, the released DNA mixture was mixed 1:1 with 60 mM sodium acetate (pH 4.6), 20 mM Zn acetate, 100 mM NaCl, and 10% glycerol ± 40 U/mL S1 nuclease (Thermo Fisher). The mixture was incubated at room temperature for 30 min before the DNA was stretched onto coverslips with a FiberComb machine (Genomic Vision).

Replication fork restart experiments were conducted with 4 × $10^5$ MDA-MB-436 cells plated in a 6-well plate and grown for 24 hr at 37°C with 5% $CO_2$. The cells were pulsed with 50 µM IdU for 30 min, briefly washed three times with prewarmed PBS and treated as indicated with DMSO, 4 mM HU, and/or 50 µM mirin for 30 min. The final concentration of DMSO used was

0.1%. The cells were then washed three times with prewarmed PBS and treated for 1 h with DMSO, 250 µM CldU, and/or 30 µM NERx 329. The final DMSO concentration was 0.6%. The cells were then washed three times with prewarmed PBS, trypsinized, and embedded in agarose plugs. Agarose plugs were lysed overnight with 2 mg/mL proteinase K (Thermo Fisher) at 50°C. The next day, the agarose plugs were washed three times with TE buffer for 1 h each at room temperature with gentle rotation. Agarose plugs were then added to 0.5 M MES pH 5.5 at 68°C for 20 min, followed by incubation at 42°C for 10 min. β-agarase (New England Biolabs) was added to digest all the agarose and release the DNA overnight at 42°C. The next day, the released DNA mixture was mixed 1:1 with 0.5 M MES, pH 5.5, before the DNA was stretched onto coverslips with a FiberComb machine (Genomic Vision).

The coverslips from both the S1 nuclease fiber combing and replication fork restart experiments were processed in the same way as described below. Coverslips were incubated at 60°C for 2 hr. DNA was denatured by incubating coverslips in 0.5 M NaOH/1 M NaCl. Coverslips were washed three times with PBS, dehydrated with an ethanol series (70%, 90%, 100%), and then air dried for 30 min. Coverslips were blocked with 3% BSA in PBST (PBS + 0.1% Tween 20) for 1 hr at 37°C. Coverslips were incubated with 1/25 mouse anti-BrdU (BD Biosciences, 347580, for IdU detection) and/or 1/50 rat anti-BrdU (Abcam, Ab6326, for CldU detection) in 3% BSA in PBST for 1 hr at 37°C. Coverslips were washed three times with PBST and incubated with 1/100 goat anti-mouse cross-adsorbed AlexaFluor488 (Invitrogen) and/or 1/100 goat anti-rat cross-adsorbed AlexaFluor594 (Invitrogen) in 3% BSA in PBST for 1 hr at 37°C. Coverslips were washed three times with PBST, mounted on slides with ProLong Diamond antifade mounting media, and allowed to cure overnight. DNA fibers were visualized with an EVOS FL Auto 2 Imaging System (Invitrogen) under a 60X oil immersion lens with GFP and TxRed filters. Individual fibers were measured via ImageJ. All fiber combing data are from duplicate experiments with a minimum of 100 fibers analyzed for each condition.

*Flow Cytometry*

MDA-MB-436 cells were plated in 12-well dishes at $2 \times 10^5$ cells/well and allowed to adhere overnight. The cells were treated with the RPAi NERx 329, the indicated PARPi, and drug combinations for 48 hours as indicated. The DMSO concentration was held constant at 0.5% for all the treatments. To analyze actively replicating cells, cultures were labeled with EdU for the final 30 minutes of the 48-hour treatment period. After treatment, the cells were washed twice with PBS, trypsinized and collected in PBS containing 1% FBS and 1 mM EDTA. After centrifugation at 400xg for 3 minutes, the cells were washed once with 1% BSA in PBS. EdU was detected with the Click-iT® EdU Flow Cytometry Assay Kit (Molecular Probes) according to the manufacturer's protocol. Briefly, the cells were fixed in Click-iT® fixative for 15 minutes at room temperature and permeabilized with Click-iT® saponin-based permeabilization and wash reagent. Click-It reactions were carried out at room temperature for 30 minutes, after which the cells were washed with Click-iT® saponin-based permeabilization and wash reagent. DNA was stained with Guava® Cell Cycle Reagent (Cytek) for 30 minutes, and data were acquired on a Guava easyCyte flow cytometer.

*FUCCI Cell Line Generation*

Twenty-four hours prior to transduction, A549 or MDA-MB-436 cells were plated at $1 \times 10^4$ cells/well in a 48-well plate. The IncuCyte Cell Cycle Green/Red Lentivirus Reagent (Sartorius, Cat# 4779) was added at an MOI of 4.5 in media containing 8 µg/mL polybrene (Millipore Sigma, Cat# TR-1003). Beginning 48 hours after viral transduction, the cells were subjected to puromycin selection at 0.6 µg/mL for 2–4 weeks and then serially diluted in 96-well plates to select single-cell clones. Puromycin-resistant clones were selected on the basis of the relatively equal intensity of the GFP and RFP markers.

*Genomic Analysis*

We followed GATK best practices[1] to process whole genome sequencing (WGS) and whole exome sequencing (WES) data for the HDH cell lines and identify short variants (SNPs and indels). FastQC (http://www.bioinformatics.babraham.ac.uk/projects/fastqc) was used to assess the quality of the sequencing reads, followed by adapter trimming with Cutadapt (https://doi.org/10.14806/ej.17.1.200 ). Clean reads were mapped to the human genome (hg19) using BWA-MEM2. PCR duplicates were removed with the command, OPTICAL_DUPLICATE_PIXEL_DISTANCE 2500. After recalibration of base quality scores with BaseRecalibrator and ApplyBQSR in GATK, variant discovery, including SNPs and INDELs, was performed with Mutect2 in tumor-only mode, using the gnomAD VCF file as a germline resource and the 1000 Genomes Project samples as the panel of normal.

### Supplemental Figures

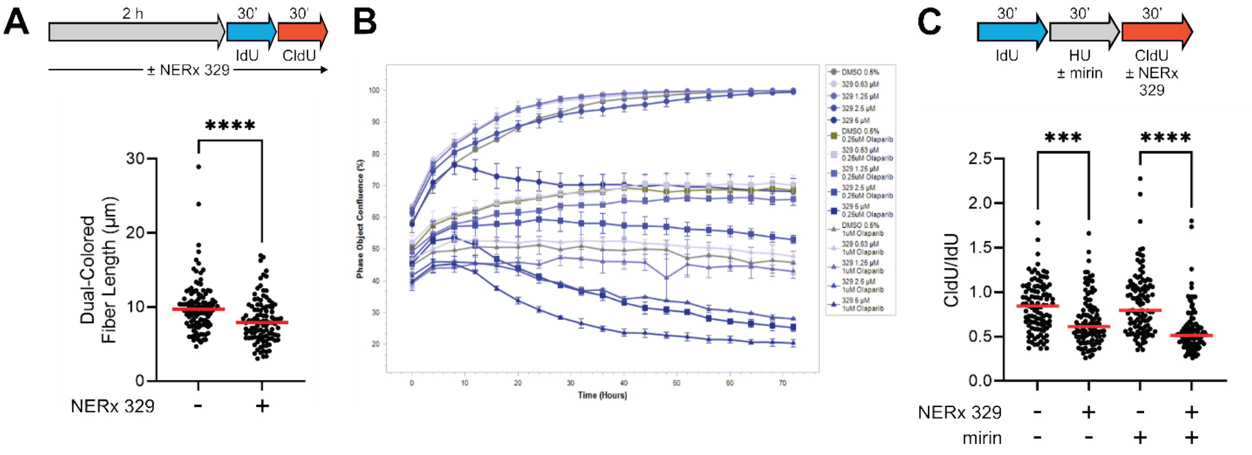


**Figure S1. A)** Labeling scheme (*top*) and quantification (*bottom*) of dual-colored fiber lengths from DNA fiber combing experiments to measure replication fork progression. A549 cells were treated with vehicle control or NERx 329 for 2 hr before labeling active replication forks with IdU for 30 min and CldU for 30 min. Median measurement is indicated by the solid red bar. **B)** Confluence of MDA-MB-436 cells pretreated for 2 days with vehicle control or olaparib, followed by the addition of vehicle control or NERx 329 (Day 0 on the graph) as monitored by Incucyte live cell imaging. Olaparib pretreatment was conducted before beginning the live cell imaging. **C)** Labeling scheme (*top*) and quantification (*bottom*) of DNA fiber combing experiments assessing replication fork restart after hydroxyurea treatment in A549 cells treated with vehicle, NERx 329, and/or mirin. Median measurement is indicated by the solid red bar.

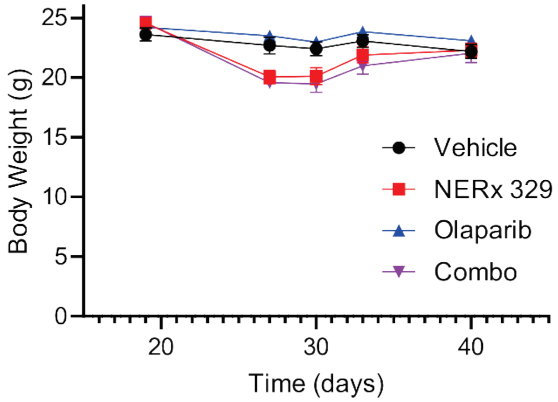


**Figure S2.** Mass of tumor-implanted NGC mice over time from the four arms of the experiment described in Figure 2.

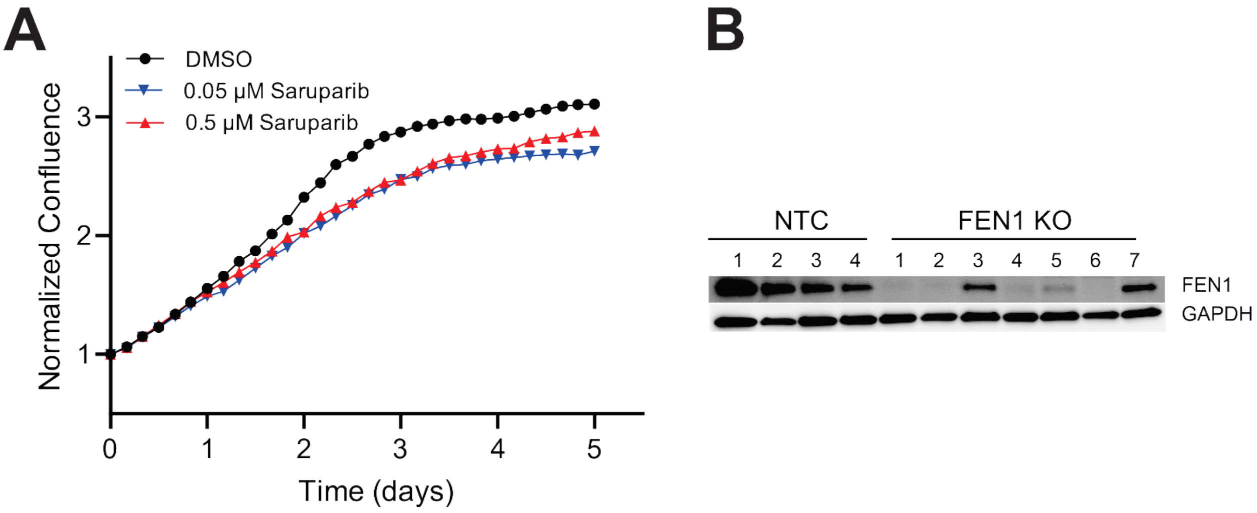


**Figure S3. A)** Confluence of MDA-MB-436 cells treated with vehicle control or saruparib as monitored by Incucyte live cell imaging. **B)** Western blot probing FEN1 expression of Cas9-expressing A549 cells treated with nontargeted control (NTC) or FEN1 gRNA.

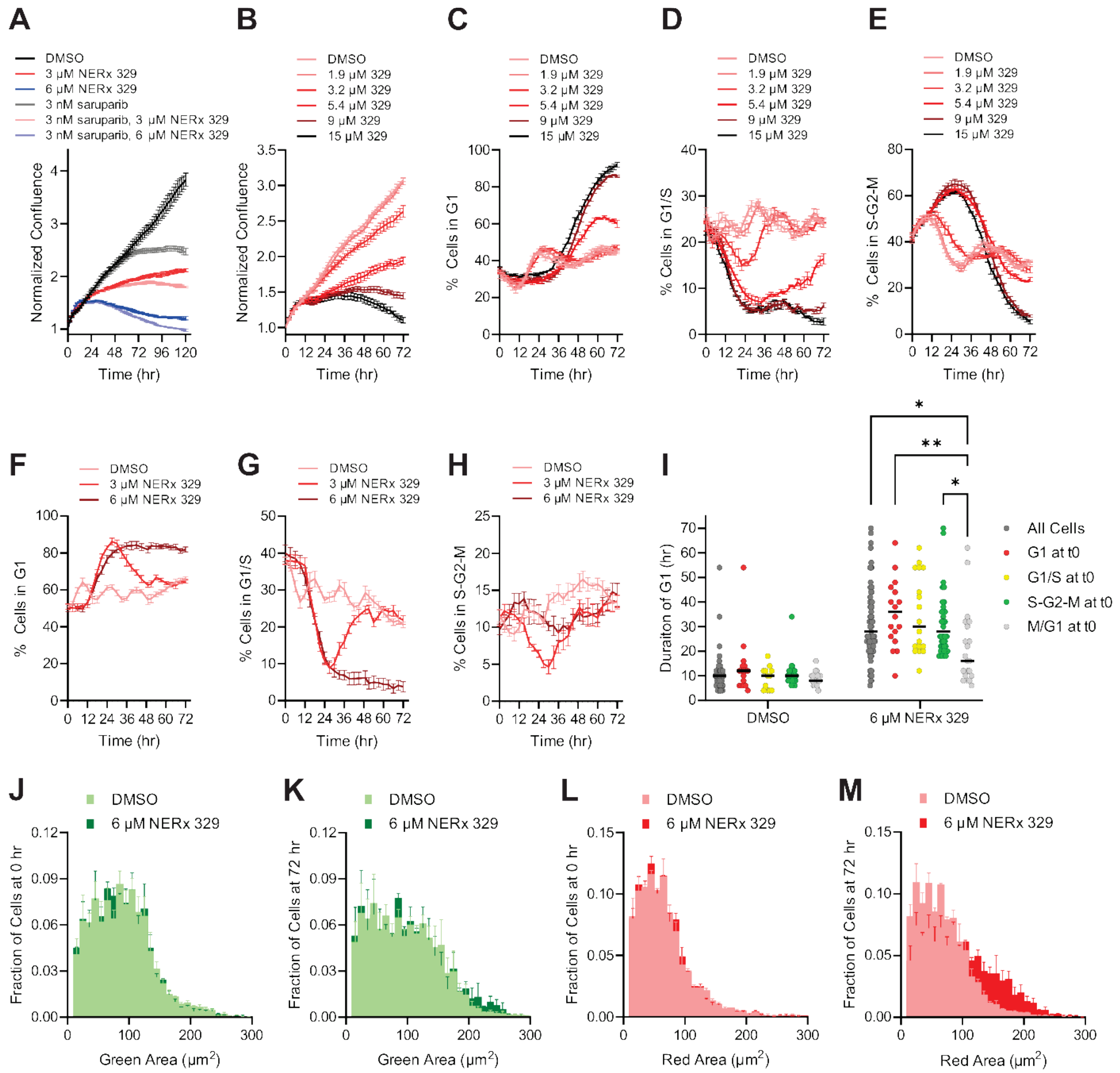


**Figure S4. A)** Confluence of MDA-MB-436 FUCCI cells treated with vehicle control, NERx 329, saruparib, or combination as monitored by Incucyte live cell imaging. **B)** Confluence of MDA-MB-436 FUCCI cells treated with vehicle control or increasing concentrations of NERx 329 as monitored by live cell imaging. Quantification of MDA-MB-436 FUCCI cells in **C)** G1, **D)** G1/S, or **E)** S-G2-M following treatment with vehicle control or increasing concentrations of NERx 329. Quantification of NSCLC A549 FUCCI cells in **F)** G1, **G)** G1/S, or **H)** S-G2-M following treatment with vehicle control or increasing concentrations of NERx 329. **I)** Quantification of MDA-MB-436 FUCCI cell G1 durations following treatment with vehicle control or NERx 329. Populations are divided according to the cell cycle phase that each cell was in upon the initiation of treatment. Area of green fluorescence at **J)** the start of treatment and **K)** after 72 hours from MDA-MB-436 FUCCI cells treated with DMSO or NERx 329. Area of red fluorescence at **L)** the start of treatment and **M)** after 72 hours from MDA-MB-436 FUCCI cells treated with DMSO or NERx 329.

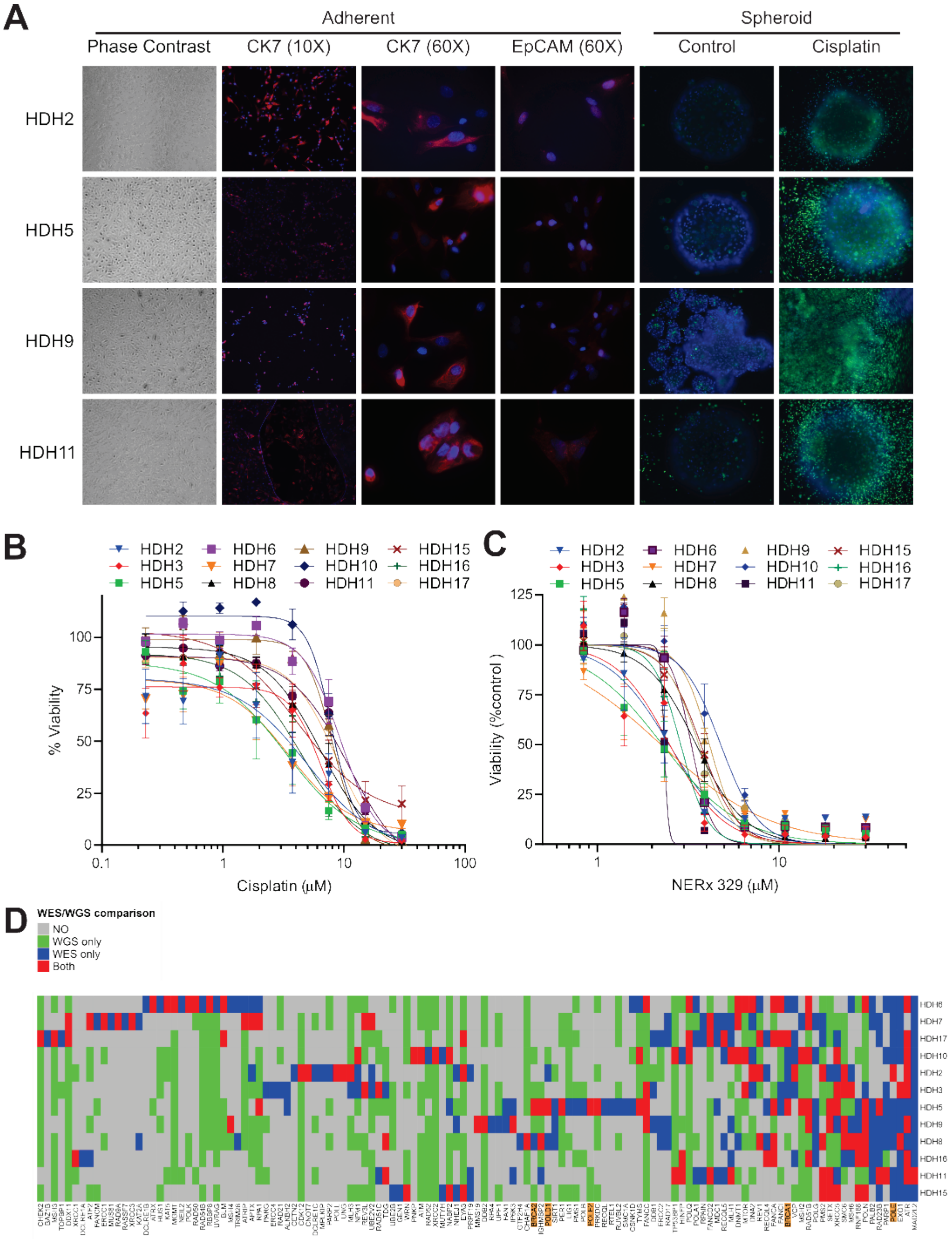


**Figure S5. A)** CK7 and EpCAM expression from adherent cultures and spheroid integrity following cisplatin treatment from representative HDH cell lines. **B)** Cell viability measurements of HDH cell lines treated with increasing concentrations of cisplatin. **C)** Cell viability measurements of HDH cell lines treated with increasing concentrations of NERx 329 normalized to vehicle control. **D)** Heat map of DDR gene mutations identified by whole exome and/or whole genome sequencing.
